# Striatal volume and functional connectivity predict weight gain in early-phase psychosis

**DOI:** 10.1101/501825

**Authors:** Philipp Homan, Miklos Argyelan, Christina L. Fales, Pamela DeRosse, Philip Szeszko, Delbert G. Robinson, Todd Lencz, Anil K. Malhotra

**Author notes:** Equal contribution (Todd Lencz, Anil K. Malhotra).

## Abstract

**Background:** Second-generation antipsychotic drugs (SGAs) are essential in the treatment of psychotic disorders, but are well-known for inducing substantial weight gain and obesity. Critically, weight gain may reduce life expectancy for up to 20-30 years in patients with psychotic disorders, and prognostic bio-markers are generally lacking. The dorsal striatum, rich in dopamine D2 receptors which are antagonized by antipsychotic medications, plays a key role in the human reward system and in appetite regulation, suggesting that altered dopamine activity in the striatal reward circuitry may be responsible for increased food craving and resultant weight gain.

**Methods:** Here, we measured striatal volume and striatal resting state functional connectivity at baseline, and weight gain over the course of 12 weeks of antipsychotic treatment in 81 patients with early-phase psychosis. We also included a sample of 58 healthy controls. Weight measurements were completed at baseline, and then weekly for 4 weeks, and every 2 weeks until week 12. We used linear mixed models to compute individual weight gain trajectories. Striatal volume and whole-brain striatal connectivity were then calculated for each subject, and used to assess the relationship between striatal structure and function and individual weight gain in multiple regression models.

**Outcomes:** Patients had similar baseline weights and body mass indices (BMI) compared to healthy controls. There was no evidence that prior drug exposure or duration of untreated psychosis correlated with baseline BMI. Higher left putamen volume and lower frontopolar connectivity predicted magnitude of weight gain in patients, and these effects multiplied when the structure-function interaction was considered.

**Interpretation:** These results provide evidence for a synergistic effect of striatal structure and function on antipsychotics-induced weight gain. Lower fronto-striatal connectivity, implicated in less optimal long-term decision making, was associated with more weight gain, and this relationship was stronger for higher compared to lower left putamen volumes.

**Funding:** Supported by NIMH grant P50MH080173 to Dr. Malhotra, grant R01MH060004 to Dr. Robinson, grant R01MH076995 to Dr. Szeszko and R21MH101746 to Drs. Robinson and Szeszko.

## Introduction

Weight gain is a major side effect of treatment with antipsychotic drugs, but relative amount of weight gain may vary considerably across patients treated with a given antipsychotic agent, especially in the first episode of illness [1].

Although all antipsychotic drugs induce weight gain, some appear to induce more than others [2, 3] and reasons for differences between drugs and between patients are not fully understood. [4] Genetic variability [5–7] and lifestyle likely play a role, but structural and functional differences in the brain’s dopaminergic reward system have not been fully understood. In addition to dopamine’s function in food reward and in the control of food intake [8] dopamine is also a key factor in schizophrenia and in antipsychotic action, suggesting that one may expect baseline differences in striatal structure and function to account for the variability in antipsychotic-induced weight gain.

The dorsal striatum, rich in dopamine D2 receptors which are antagonized by antipsychotic medications, plays a key role in the human reward system and in appetite regulation, suggesting that altered dopamine activity in the striatal reward circuitry may be responsible for increased food craving and resultant weight gain [9]. In line with this notion, previous work has shown that decreased baseline functional activity in the putamen predicted amount of future antipsychotic weight gain [10] and olanzapine-induced activity in the dorsal striatum was associated with excessive eating behavior [11]. Supporting the hypothesis of altered reward processing in schizophrenia, these results suggest that decreased reward anticipation in the putamen before antipsychotic treatment may predispose to weight gain under antipsychotic treatment. However, we are not aware of studies in weight gain using risperidone and aripiprazole, two widely used SGAs [12]. Furthermore, in addition to baseline function, variability in striatal structure and functional connectivity may also contribute to weight gain. Indeed, previous studies have shown higher striatal volumes in addictive behavior [13, 14] as well as weaker fronto-striatal connectivity in obesity [15]. Thus, we measured striatal volume and striatal resting state functional connectivity in patients with early-phase psychosis at baseline, and weight gain over the course of 12 weeks of treatment with risperidone and aripiprazole, and hypothesized that higher striatal volume and lower cortico-striatal connectivity would be associated with more weight gain [15, 16].

## Materials and Methods

### Participants

We used two early-phase psychosis cohorts from two separate 12-week clinical trials with a similar design and similar treatment effects. Details have been published previously [17] and are summarized in Table 1. Written informed consent was obtained from adult participants and the legal guardians of participants younger than 18 years. All participants under the age of 18 provided written informed assent. The study was approved by the Institutional Review Board (IRB) of Northwell Health. We also included a sample of 58 age-matched healthy controls (Table 1).

**Table 1:** Sample characteristics. *Abbreviations*: a, significant difference between patients and controls (*P* < 0.05) after adjusting for age, sex, age-by-sex, body mass index and total intracranial volume; b, significant predictor of weight gain in schizophrenia (*P* < 0.05); NOS, not otherwise specified; PSY, early-phase psychosis patients; HC, healthy controls; BPRS, Brief Psychiatric Rating Scale; BMI, body mass index; DUP, duration of untreated psychosis.

| Characteristic | PSY | Mean | SD | HC | Mean | SD |  |
| --- | --- | --- | --- | --- | --- | --- | --- |
| Males | 58 |  |  | 25 |  |  |  |
| Females | 23 |  |  | 33 |  |  |  |
| Age, years | 81 | 21.5 | 5.5 | 58 | 28.1 | 11.9 |  |
| Schizophrenia | 57 |  |  |  |  |  |  |
| Schizophreniform disorder | 14 |  |  |  |  |  |  |
| Psychotic disorder NOS | 5 |  |  |  |  |  |  |
| Schizoaffective disorder | 2 |  |  |  |  |  |  |
| Bipolar I disorder (with psychotic features) | 3 |  |  |  |  |  |  |
| Prior medication exposure | 74 | 8.4 | 29.6 |  |  |  |  |
| Medication naive | 28 |  |  |  |  |  |  |
| Medication 2 weeks or less | 70 |  |  |  |  |  |  |
| Education, years | 76 | 12.2 | 2.2 | 58 | 13.7 | 2.9 |  |
| Weight, lb | 81 | 155.1 | 35.2 | 58 | 156.1 | 28.6 |  |
| BMI | 81 | 23.6 | 4.4 | 57 | 24.7 | 4.1 |  |
| BPRS total | 81 | 42.7 | 7.5 |  |  |  |  |
| DUP, weeks | 76 | 111.9 | 177.1 |  |  |  |  |
| L Caudate, mm<sup>3</sup> | 81 | 4061.7 | 552.4 | 58 | 3922.5 | 590.6 |  |
| L Hippocampus, mm<sup>3</sup> | 81 | 4032.7 | 383 | 58 | 4107.4 | 455.8 |  |
| L Putamen, mm<sup>3</sup> | 81 | 6381.8 | 768.1 | 58 | 6316.2 | 843.6 | b |
| R Caudate, mm<sup>3</sup> | 81 | 4223.4 | 526.3 | 58 | 4103.5 | 588 |  |
| R Hippocampus, mm<sup>3</sup> | 81 | 4086.9 | 403.9 | 58 | 4201 | 416.2 | a |
| R Putamen, mm<sup>3</sup> | 81 | 6187.1 | 766.9 | 58 | 6042.7 | 850.4 |  |

### Weight assessments and analysis

Weight measurements were completed at baseline, and then weekly for 4 weeks, and every 2 weeks until week 12. To estimate the individual level of weight gain during this time frame, we used linear mixed models which allowed us to compute individual weight gain trajectories [18]. Specifically, we used weight as the dependent variable and time (measured in days from baseline) as a continuous predictor, and included a random intercept as well as a random slope for day. The random slope for each participant is an index of the individual weight gain and was the measure of primary interest in this study. Random slopes were estimated using Restricted Maximum Likelihood (REML) and used as the dependent variables in our structural and functional imaging analysis (see below). All analyses were conducted in R version 3.5.1 (2018-07-02). Data and code of the current study are available online at http://github.com/philipphoman/bmi.

### Structural imaging and analysis

Magnetic resonance imaging exams were conducted on a 3-T scanner (GE Signa HDx). We acquired anatomical scans in the coronal plane using an inversion-recovery prepared 3D fast spoiled gradient (IR-FSPGR) sequence (TR = 7.5 ms, TE = 3 ms, TI = 650 ms, matrix = 256 × 256, FOV = 240 mm) which produced 216 contiguous images (slice thickness = 1 mm) through the whole brain. After image processing and segmentation with Freesurfer 5.1.0, we measured the volumes of putamen and caudate and also included the hippocampus as control region.

We then computed multivariable regressions for each region of interest, entering the individual weight gain slopes as the dependent variable and a predictor for subcortical volume. To adjust for unspecific confounders, we included additional variables in these models, namely age, sex, the age-by-sex interaction, baseline body mass index (BMI), duration of untreated psychosis (DUP), and total intracranial volume.

### Functional imaging and analysis

As described previously [17] we obtained resting-state functional scans during a session of 5 min in duration. The resting state scan included 150 echo-planar imaging (EPI) volumes with a TR = 2000 ms, TE = 30 ms, matrix = 64 × 64, FOV = 240 mm, slice thickness = 3 mm, and 40 continuous axial oblique slices (one voxel = 3.75 × 3.75 × 3 mm). During resting state scanning, participants were asked to close their eyes and instructed not to think of anything in particular. We used FSL (http://www.fmrib.ox.ac.uk) for preprocessing of the resting-state scans. After discarding the first four EPI volumes, each participant’s functional image was registered to a corresponding structural T1 image using a linear transformation with six degrees of freedom. This structural image was normalized by a 12-parameter affine transformation to MNI-152 space. The combination of these transformations was then applied to each individual’s functional dataset. Rigid body motion correction was performed with MCFLIRT and skull stripping was performed with BET.

Images were spatially smoothed with a 5-mm FWHM Gaussian kernel. The resulting time series was then high pass filtered at 0.01 Hz. White matter (WM) and cerebrospinal fluid (CSF) masks were generated using FAST by segmentation of each individual’s structural image. Mean signal within these masks was extracted. For removal of nuisance variables, each individual’s 4D time series the data were regressed with eight predictors in a general linear model: WM, CSF, and six motion parameters. To avoid interference with our connectivity measures, the global mean was not included in this calculation.

Since our structural analysis revealed a single region to be predictive of weight gain, i.e., the left putamen, we used this as a seed region in our functional connectivity analysis. Adopting the approach by Di Martino and colleagues [19] 4 × 4 × 4 mm spheres were defined in the subregions of the left putamen, including the dorsal rostral putamen (x = −25, y = 8, z = 6), dorsal caudal putamen (x = −28, y = 1, z = 3), and ventral rostral putamen (x = −20, y = 12, z = −3). We computed correlation maps for each participant for all 3 of our ROIs by extracting mean activity time courses from each seed region, and by calculating whole-brain voxel-wise correlation maps with the extracted waveform as a reference. The resulting correlation maps were *z*-transformed.

Mean frame-wise displacement (FD) was calculated in FSL and was also included to control for the residual effect of head motion. Given that use of the data scrubbing to eliminate motion-related artifact offers little advantage over group-level corrections and can correct the data incompletely [20] we accounted for head motion at the group-level by including mean FD as a nuisance covariate [20, 21]. Group level analyses were performed independently for each ROI in FSL’s FLAME. For each ROI, all maps were entered into a general linear model with age, sex, FD, and baseline BMI as covariates, and the individual weight gain slope as outcome measure. Significance was defined voxel-wise at *z* > 3.1, with cluster correction at *P* < 0.05.

## Results

Patients and controls had similar weights and body mass indices (BMIs) at baseline (that were also within the normal range; Table 1). There was no evidence that the duration of untreated psychosis correlated with BMI (*β* = −0.01, *t* (62) = −0.14, *P* = 0.889) or that prior exposure to antipsychotics (before entering the trial) was correlated with baseline BMI (*β* = 0.19, *t* (68) = 1.59, *P* = 0.117). Furthermore, non drug-naive patients did not have a higher baseline BMI (M = 23.58, SD = 4.49) compared to healthy controls (M = 24.68, SD = 4.08). The left hippocampal volume was significantly reduced in patients compared to controls (Table 1), but no significant case-control differences were observed for any of the striatal regions.

The left putamen volume predicted magnitude of weight gain (Fig. 1a) in patients (*β* = 0.31, *t* (68) = 2.18, *P* = 0.032; Fig. 1b), with larger baseline volumes associated with greater weight gain during the subsequent trial.

**Figure 1:**
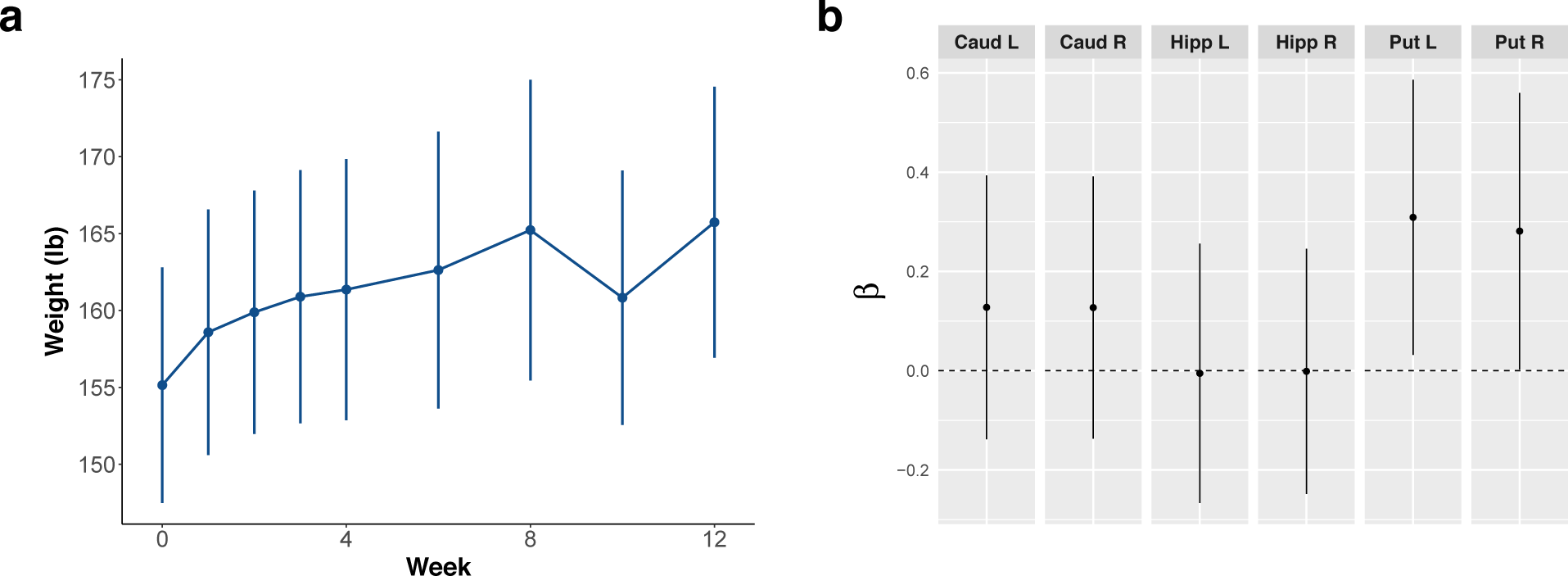
Striatal volume predicts weight gain in early-phase psychosis patients (*N* = 81). **a. Weight increased significantly over the course of the trial.** Means with 95% confidence intervals are shown. **b. Weight gain in patients was predicted specifically by the left putamen volume.** Standardized beta coefficients with 95% confidence intervals are shown. All models were adjusted for age, sex, age-by-sex, duration of untreated psychosis, intracranial volume, and body mass index, none of which were significant predictors of weight gain. Error bars not touching the zero line indicate significant effects (*P* < 0.05). *Abbreviations*: Caud, caudate; Hipp, hippocampus; Put, putamen; L, left, R, right.

Following up on the left putamen volume finding, we tested for functional resting-state connectivity [17] between the left putamen and the whole brain that was associated with weight gain in 75 patients. For the left dorsal rostral putamen, we found that decreased functional connectivity with the left medio-lateral frontal pole also predicted amount of weight gain (Fig. 2a, b). Left putamen volume and fronto-striatal connectivity were not significantly correlated (*β* = −0.15, *t* (73) = −1.31, *P* = 0.194).

**Figure 2:**
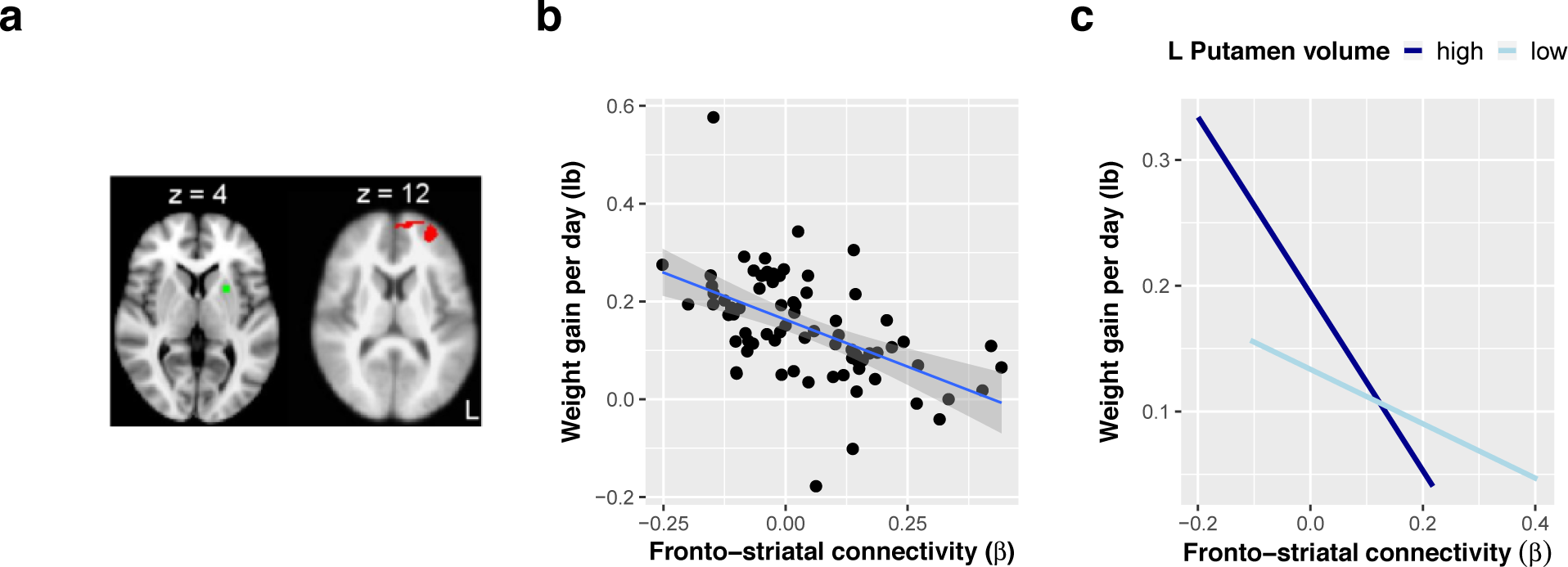
Fronto-striatal connectivity and the interaction of striatal structure and function predict weight gain in early psychosis patients (*N* = 75). **a, b. Lower functional connectivity between the left putamen and the left frontal pole predicted weight gain.** Following up on the left putamen volume finding, we tested for functional connectivity between the left putamen and the whole brain that was associated with weight gain. The left dorsal rostral putamen (green) was used as seed region, and the left frontal pole (red) was the only connected region surviving a voxel level threshold of *z* > 3.1 and a cluster threshold of *P* < 0.05. **c. Interaction of striatal structure and function in predicting weight gain.** To illustrate the significant interaction between left putamen volume and connectivity, we plotted regression lines for weight gain and connectivity for the highest quartile (25%) of putamen volume and the lowest quartile (25%) of putamen volume.

Notably, the regression models were adjusted for age, sex, intracranial volume, baseline BMI, and duration of untreated psychosis; none of these variables showed a significant association with weight gain. In addition, to verify that the results were robust to the type of medication administered (risperidone vs. aripiprazole), we extended the models by including interactions of medication type and volume as well as medication type and connectivity, respectively, in the two regression models. There was no evidence for an effect of medication type on the weight gain and volume association (*β* = 0.01, *t* (66) = 0.11, *P* = 0.914) or the weight gain and connectivity association (*β* = 0.06, *t* (71) = 0.54, *P* = 0.594). In addition, no interactions with volume or connectivity were found for prior exposure to antipsychotics and medication dose during the trial.

These results suggest that patients with higher striatal volume and more negative fronto-striatal connectivity gained more weight during the clinical trial. To investigate a potential synergistic effect of structure and function, we added functional connectivity indices as well as the functional connectivity by volume interaction to an extended regression model. This model allowed us to test whether striatal volume moderated the association between fronto-striatal connectivity and weight gain. Indeed, we found evidence for an interaction of volume and connectivity (*β* = −0.38, *t* (65) = −3.36, *P* = 0.001; Fig. 2c), indicating that the negative relationship between fronto-striatal connectivity and weight gain was weaker with lower compared to higher striatal volumes. Note that this effect remained significant after excluding patients who had bipolar I disorder and a recent manic episode with psychotic features (*N* = 3); and after adjusting the model for type of medication, medication dose during the trial, and prior exposure to antipsychotic drugs. However, since the functional connectivity measure that entered this model is not an independent measure of the effect size of the fronto-striatal connectivity [22] we also calculated a leave-one-out cross validation to derive an independent measure of the effect size of this interaction, and found that this effect remained significant (*r* (65) = 0.3, *P* = 0.013).

## Discussion

Here we showed that striatal volume and fronto-striatal connectivity predicted degree of weight gain associated with antipsychotic treatment. Lower connectivity between the left frontal pole and the left putamen was associated with more weight gain, and this relationship was stronger for higher compared to lower left putamen volume. In accordance with previous studies, we also found lower hippocampal volumes in patients compared to controls [23–25].

We focused our study on the striatum, in line with this region’s key role in reward processing and weight gain. Previous studies have shown that treatment with antipsychotics in healthy controls induced an increase in reward activation in the dorsal striatum that was correlated with excessive eating [11] and that attenuated reward anticipation normalizes under treatment with antipsychotics in patients with schizophrenia [26, 27]. However, although altered reward processing is likely to play a role [10], antipsychotics-induced weight gain remains a complex issue [4]. First, one could ask why patients did not gain weight before the antipsychotic treatment if baseline alterations in reward processing are the main cause for weight gain. Indeed, there is evidence suggesting that patients might show metabolic aberrations already before they start treatment, possibly due to unhealthy lifestyle, illness neurobiology, and genetic factors [28–31]. Furthermore, it is important to consider that some patients in the current study were not naive to antipsychotic drugs but had been exposed to antipsychotic treatment prior to inclusion. Thus, weight gain may have occurred during this time frame. Nevertheless, we did not find evidence that baseline BMI correlated with days of prior drug exposure or that non drug-naive patients had a higher baseline BMI compared to controls, speaking against substantial weight gain before the trial. The current trial of 12 weeks, although short, thus appears long enough to induce substantial weight gain with considerable inter-subject variability, especially in subjects with minimal prior antipsychotic exposure [1]. Second, one might ask how antipsychotics and some of their counterparts, dopaminergic agonists, can both induce weight gain, as shown for Parkinson disease [32]. A possible explanation is that weight gain in Parkinson disease is triggered mainly by compulsive eating [32, 33] whereas a decreased reward experience may be the main cause for weight gain induced by antipsychotics. It is also noteworthy that increased dopamine transmission through sympatomimetics such as amphetamine typically results in suppressed appetite.

A probable mechanism for weight gain is thus via altered reward processing in the striatum. The striatum is part of the brain’s reward circuitry which has well-known dopaminergic components as well as neurotransmitters including opioids, serotonin, and cannabinoids [8, 11, 34–36]. Accordingly, food consumption increased striatal dopamine in a previous study using positron emission tomography (PET) and this increase was positively correlated with meal pleasantness [35]. In addition, food cues elicited dorsal striatum dopamine increase and this increase correlated with hunger perception and food desire [37]. In obesity, dopamine has also been implicated in food reward and the control of eating behavior [38–40]. More specifically, functional magnetic resonance imaging studies in obesity have found altered activation in dopaminergic reward-related areas. Although these studies have described enhanced striatal activation in response to food-related cues [41–44] the actual consumption of food elicited less activation in these areas [42, 43]. As previously suggested [11] these patterns are consistent with an imbalance between (increased) reward expectation and (decreased) reward experience, most likely through dopaminergic disruptions. Overeating could then be seen as a compensation to obtain the anticipated exaggerated reward [11].

Our study extends this model by considering the cortico-striatal functional connectivity which has been implicated in long term, inference-based decision making that can be biased by short-term reward experiences [45]. Accordingly, we found that the strength of this connectivity was correlated with less weight gain. Furthermore, we found that the association with weight gain was moderated by left putamen volume, with higher volumes together with lower connectivity predicting more weight gain. The moderating role of the putamen volume suggests a synergistic effect of striatal structure and function, with increased putamen volumes, possibly due to reduced endogenous dopamine availability in the striatum [13, 14] multiplying the effect of lower cortico-striatal connectivity on weight gain [15, 16].

Some limitations merit comment. First, although the effect size of the striatal structure-function and weight gain relationship was medium (*r* = 0.3), these data may provide the first biomarker-based tool for identification of patients at very high risk for weight gain upon initiation of antipsychotic treatment. Furthermore, we used two different SGAs in this study, namely risperidone and aripiprazole. Although it is important to underscore that all current antipsychotics share affinity for the dopamine D2 receptors, they may still differ from a pharmacological point of view. For example, risperidone and aripiprazole share D2 receptor antagonism to induce an antipsychotic effect [46] but different receptor systems may be involved in side effects such as weight gain. Apart from the striatum [10] weight gain has been associated with cortical 5-HT2A receptors in quetiapine monotherapy [47]. In addition, olanzapine exposure in healthy controls indicated negative effects on the peripheral metabolism. These findings suggest that antipsychotic induced weight gain may involve additional central and peripheral aspects apart from the striatum. However, in the current study we did not find evidence that the type of medication interacted with the effects of striatal volume or fronto-striatal connectivity, suggesting similar effects of both drugs, which is also consistent with our previous finding for a larger trial [12] where no significant differences on weight gain were found between aripiprazole and risperidone. In addition, weight gain differences between aripiprazole and risperidone found in previous studies might reflect a difference in the sedative effect, with risperidone causing more sedation than aripiprazole [2]. Finally, future studies with repeated measurements of structural and functional imaging should investigate whether our findings reflect indeed a trait marker in patients prone to weight gain.

In conclusion, the current study showed a synergistic effect of striatal structure and function on antipsychotics-induced weight gain. Lower fronto-striatal connectivity, implicated in unfavorable long-term decision making, was associated with more weight gain, and this relationship was stronger for higher compared to lower left putamen volumes. This suggests that an imaging marker at baseline may identify patients who are prone to substantial weight gain during treatment.

## Acknowledgments

Supported by NIMH grant P50MH080173 to Dr. Malhotra, grant R01MH060004 to Dr. Robinson, grant R01MH076995 to Dr. Szeszko and R21MH101746 to Drs. Robinson and Szeszko. The authors thank Dr. Lauren Hanna and Dr. Juan Gallego for their careful clinical oversight of the study. They acknowledge their patients, their patients’ families, and their psychiatry research support staff.

## Conflict of Interest

Dr. Robinson has been a consultant to Costello Medical Consulting, Innovative Science Solutions, Janssen, Lundbeck, Otsuka and US WorldMeds and has received research support from Otsuka. Dr. Malhotra has served as a consultant for Forum Pharmaceuticals and has served on a scientific advisory board for Genomind. Dr. Lencz is a consultant for Genomind. The other authors report no financial relationships with commercial interests.

